# Multi-omics insights into quinolizidine alkaloid biosynthetic architecture in narrow-leafed lupin genotypes with contrasting alkaloid regulation

**DOI:** 10.64898/2026.01.08.698079

**Authors:** Katarzyna Czepiel, Aleksandra Burdzińska, Karolina Susek, Agnieszka Kiełbowicz-Matuk, Grzegorz Koczyk, Magdalena Kroc

**Author notes:** Correspondence (M.K.). (K.C.); (K.S.), (A.B.), (A.K.M.), (G.K.).

## Abstract

Quinolizidine alkaloids restrict the use of lupin seeds as food and feed. In narrow-leafed lupin, low-alkaloid content in most cultivars has been traced to the recessive *iucundus* locus, with the RAP2-7 transcription factor as a candidate regulator, yet the molecular basis of alternative low-alkaloid sources remains unclear. Here we provide a comparative view of alkaloid pathway and its regulation across two genetic backgrounds, the *iucundus* and *Iucundus* contrast and the Bryansk low-alkaloid background carrying the *Iucundus*-type RAP2-7 allele. This multi-omic framework integrates alkaloid profiling and transcriptomics alongside RAP2-7 DNA binding characterization, and sequence-level motif and variant analyses. Alkaloid profiles revealed genotype-specific differences, with Bryansk lines showing a distinct, sparteine-enriched and lupanine-depleted chemotype relative to *iucundus* and *Iucundus*. Using *Iucundus* line as a reference, transcriptome analyses highlighted candidate genes associated with low-alkaloid *iucundus* and Bryansk backgrounds, spanning enzymes, transporters and putative regulators. Consistent with a key role of RAP2-7, DAP-seq summits in *Iucundus* and Bryansk contained a clear AP2-like motif, whereas the *iucundus* background showed both strong depletion of high-confidence peaks and no defined motif. *In silico* modelling of RAP2-7 bound to its DNA motif, combined with DAP-seq and expression data, supported reduced binding of the *iucundus* variant relative to *Iucundus,* as well as a crucial mutation within the promoter of key acyltransferase (*LaAT*). Collectively, these data extend lupin transcriptomic resources and refine models of alkaloid biosynthesis beyond classical *iucundus* sources, and within this comparative framework, provide the first comprehensive molecular characterization of Bryansk low-alkaloid lines.

## Introduction

Plants synthesize a diverse spectrum of specialized metabolites that often contribute to their ecological fitness and adaptive potential. In lupins, quinolizidine alkaloids (QAs) form a major class of such compounds, underpinning plant defense and shaping end-use quality of seeds [1]. Given their toxicity, feed safety requires QA level below ∼0.02% of seed dry weight, a threshold used to designate sweet genotypes [2].

A comprehensive understanding of a crop trait requires elucidation of its molecular basis, including the underlying pathway, regulation and transport, together with characterization of genetic resources to capture allelic diversity and relevant breeding sources. In lupins, studies of QA biosynthesis have mapped parts of the pathway, including known enzymes and their organ specific expression patterns, yet the pathway as a whole remains not yet fully elucidated [3–15]. Moreover, seed QA profiles in lupin germplasm have been partly characterized, revealing variation relevant to breeding [16–20]. Here, we focus on narrow leafed lupin (*Lupinus angustifolius* L., NLL), where low seed QA may derive from several genetic factors [16]. Among these, the recessive *iucundus* locus is the most common and best characterized, with *RAP2-7* proposed as its key regulatory factor. Additionally, in the NLL collection, an R196S substitution in exon 4 of RAP2-7 was strongly correlated with reduced seed QA content and therefore likely reflects a partial reduction in RAP2-7 activity [11,12]. Interestingly, the three Bryansk lines harbor an alternative low seed QA locus distinct from *iucundus*. Their leaves display high expression of known QA-related genes, and they carry the *RAP2-7* allele associated with high-alkaloid phenotype [11,12]. A comprehensive understanding of the QA pathway in NLL will therefore require analyses that address the role of *iucundus* and consider additional genetic and regulatory factors leading to low QA seeds.

Plant alkaloid research increasingly uses complementary datasets that combine organ resolved alkaloid and expression profiles with genome assemblies, single cell datasets, and long read resources that refine gene model. Such integrative strategies have supported pathway reconstruction, enzyme and transport assignment, gene cluster detection, and candidate biosynthetic genes selection in several systems, i.e. for monoterpenoid indole alkaloids (MIA) in *Catharanthus roseus* L. [21–25] and benzylisoquinoline alkaloids (BIA) in opium poppy [26–28]. To fully interpret plant alkaloid pathways, alongside biosynthetic reconstruction, the regulatory layers need to be resolved. DNA affinity purification sequencing (DAP-seq) is a promising approach that enables genome wide *in vitro* mapping of transcription factor binding sites (TFBS) and serves as an antibody-free alternative to ChIP-seq [29,30]. This approach, novel in lupin, enables mapping of regulatory networks for key agronomic traits, including alkaloids.

In this work we investigate two QA regulatory models in NLL: the *iucundus* locus contrast, represented by 83A:476 (*iuc*, low-alkaloid) and P27255 (*Iuc*, high-alkaloid), and an alternative low-alkaloid regulatory type represented by a set of Bryansk lines (95826, 95927 and 95928) (see Materials and Methods for details). Across these backgrounds, we profiled QA content, composition, and the expression of pathway genes across multiple organs, to resolve their genetic and regulatory contrasts. Complementary long read and short read RNA sequencing of leaves and stems refined gene annotations and revealed genotype specific expression differences. Finally, allele specific RAP2-7 binding patterns across the NLL genome in the two regulatory backgrounds were characterized using DAP-seq. Notably, this work provides the first integrated molecular characterization of the Bryansk regulatory type. Together, these datasets uncover the multilayered and genotype specific nature of QA biosynthesis in lupin, provide an integrated framework to compare established and alternative mechanisms leading to low seed QA and to prioritize candidates for functional validation.

## Results

### 1. Alkaloid quantification in lupin organs

#### Total alkaloid content (TAC)

TAC varied strongly between organs and genotypes (Table 1). Bryansk lines exhibited markedly lower TAC than P27255, but higher than 83A:476. Differences were greatest in leaves, with the Bryansk lines containing 10–25× more alkaloids than 83A:476, while remaining well below P27255 (Table 1). A similar pattern was observed in stems, flowers and pods, with TAC above 1% in *Iucundus* and below 0.1% in the Bryansk lines. TAC increased from vegetative to reproductive organs (except pods), reaching the highest values in seeds. Notably, the Bryansk lines exceeded 83A:476 by three to five-fold in green seeds, while in dry seeds both were comparable and well below P27255.

**Table 1.** Total alkaloid content (TAC, % dry weight) across organs of NLL in *iucundus* (83A:476), *Iucundus* (P27255), and Bryansk lines (95826, 95927, 95928).

| Organ* | TAC (% of dry weight) |  |  |  |  |
| --- | --- | --- | --- | --- | --- |
|  | 83A:476 | P27255 | 95826 | 95927 | 95928 |
| Leaves | 0.0017 | 0.7790 | 0.0448 | 0.0178 | 0.0198 |
| Stems | 0.0298 | 1.1219 | 0.0582 | 0.0452 | 0.0391 |
| Flowers | 0.0176 | 3.7059 | 0.0884 | 0.0946 | 0.0443 |
| Green pods | 0.0112 | 1.0373 | 0.0297 | 0.0230 | 0.0224 |
| Green seeds | 0.0333 | 2.9495 | 0.1585 | 0.1841 | 0.1148 |
| Dry seeds | 0.0648 | 2.3891 | 0.1354 | 0.0905 | 0.0746 |

\* Organs as defined in Materials and Methods (Table 4).

#### Distribution of individual alkaloids across plant organs

Qualitative QA composition varied by genotype and organ, with the Bryansk lines showing broadly similar profiles (Fig. 1; Table S1). Lupanine dominated in P27255 (∼80%), and was moderate in 83A:476 and the Bryansk lines (∼30–40%). 13-Hydroxylupanine was slightly higher in the Bryansk lines and 83A:476 (∼25–30%) than in P27255 (∼15%) and, in the Bryansk lines, showed an accumulation trend opposite to the other genotypes, increasing from leaves to seeds. Isolupanine was most abundant in the Bryansk lines (16–19%) and lowest in P27255 (3%), typically decreasing toward seeds. Angustifoline was inconsistently detected, and most abundant in dry seeds of the Bryansk lines and 83A:476. Notably, the Bryansk lines accumulated much more sparteine (∼20%) than P27255 and 83A:476 (∼0.5%), except in 83A:476 pods. Other alkaloids were low and infrequent, with P27255 displaying the broadest spectrum consistent with its high TAC, whereas 83A:476 distinguished by substantial multiflorine accumulation in stems and seeds.

**Figure 1.**
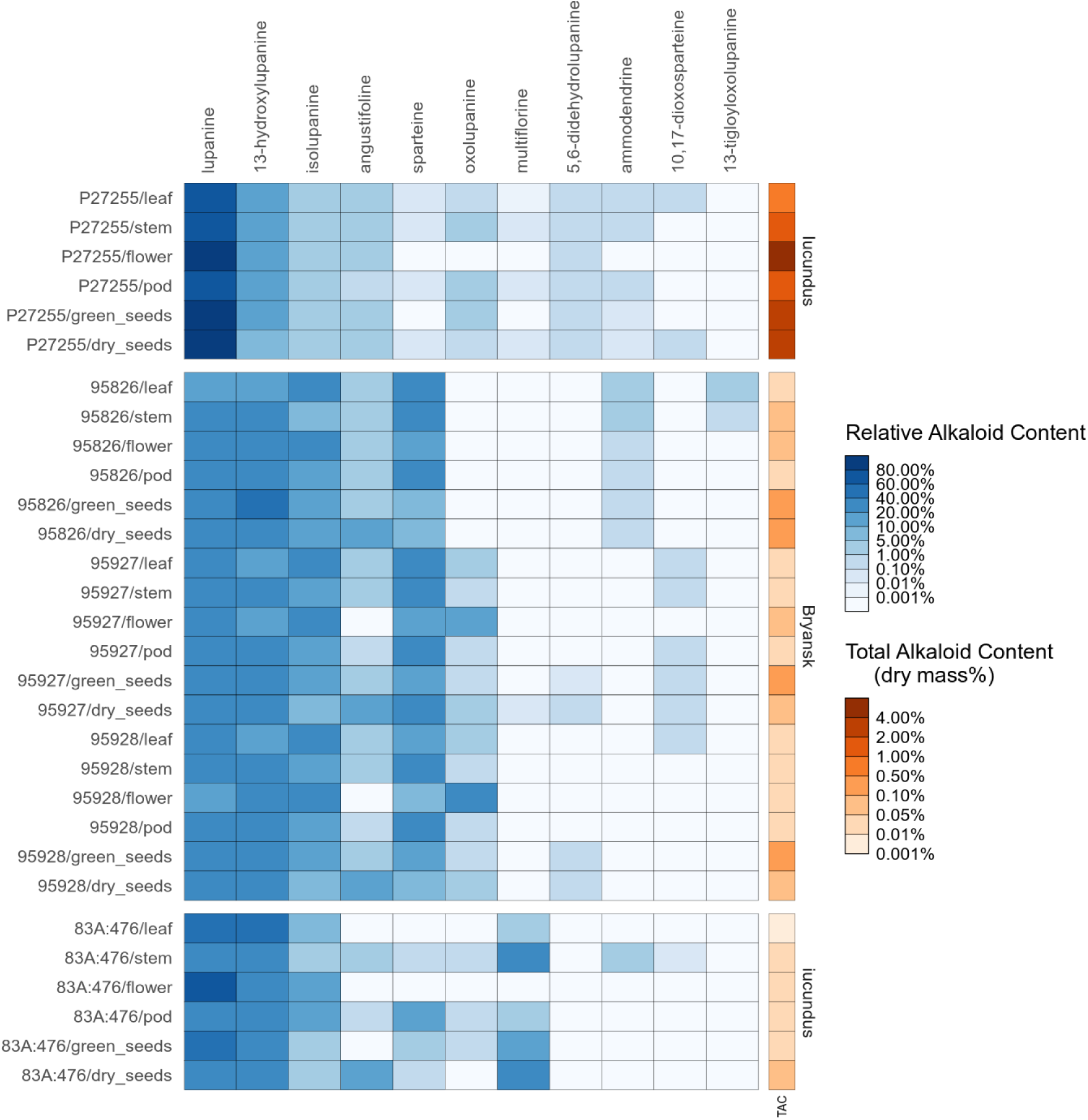
Heatmap of organ resolved QA profiles in NLL assessed by gas chromatography. Each cell shows the percent contribution of a compound to total alkaloids in a sample. The adjacent bar indicates total alkaloid content (percent dry mass). Values are means of three biological replicates.

### 2. Organ specific expression of QA-related genes across regulatory types

Across all genes examined, expression differences between P27255 and 83A:476 references were most evident in leaves and pods (Fig. 2). Stems and flowers did not show clear or consistent separation between the two genotypes, while expression in seeds was low or variable. In the Bryansk lines expression of the assayed genes was comparable to that observed in P27255. Overall, in P27255 and the Bryansk lines, expression of the assayed genes was most pronounced in green aerial lupin organs (leaves, stem and pods), with *LDC* showing the highest and *RAP2-7* generally the lowest relative expression among the analyzed genes.

**Figure 2.**
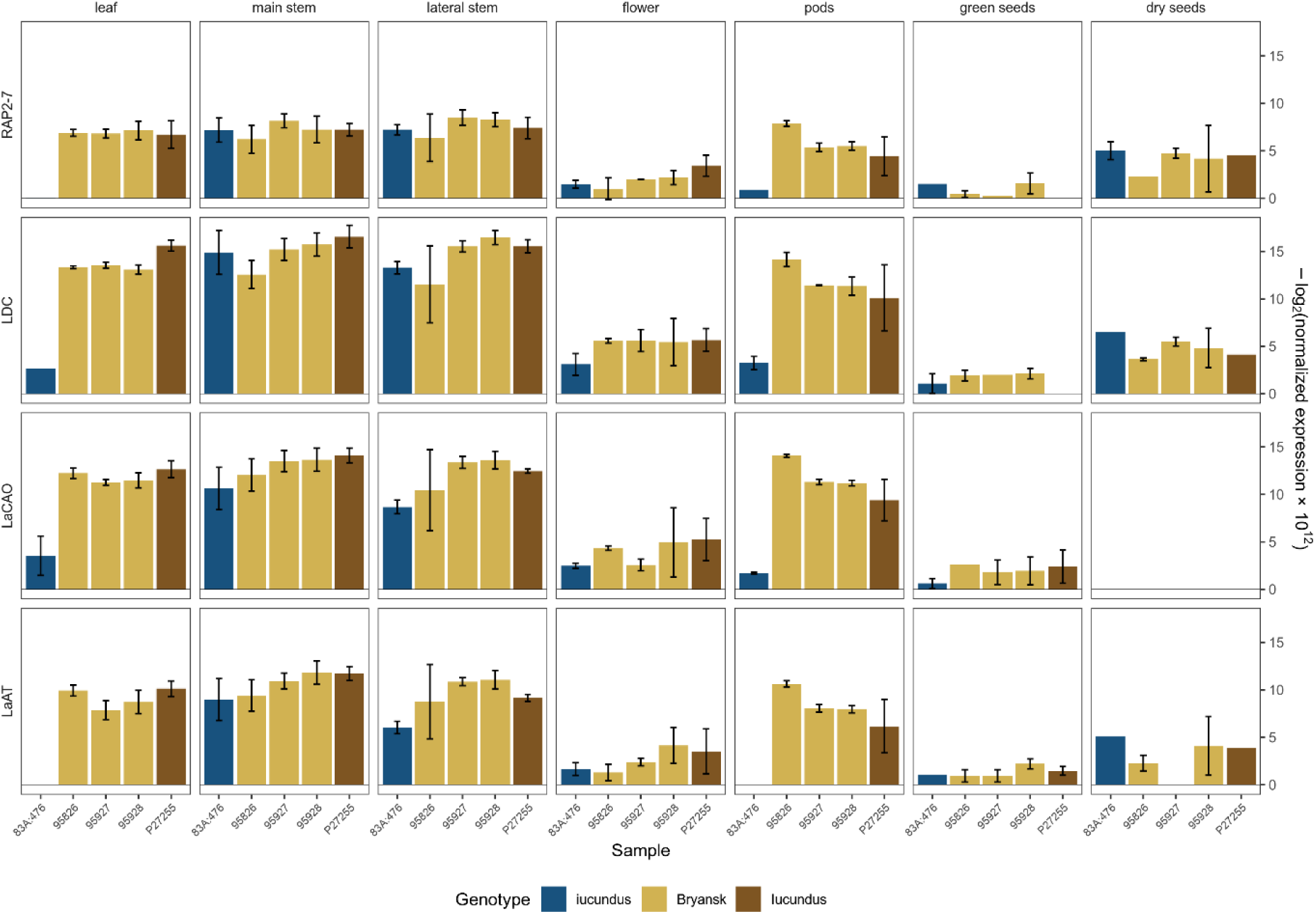
Organ specific expression of genes involved in QA pathway in five NLL genotypes, assessed by qPCR. Bar plots show mean normalized expression with standard error. Bar colors indicate genotypes grouped as *iucundus*, Bryansk lines and *Iucundus* as shown in the legend.

### 3. NLL transcriptome analysis using PacBio Iso-Seq and RNA-Seq

Comprehensive NLL transcriptome characterization was achieved by integrating PacBio Iso-seq and Illumina RNA-seq for leaf and stem of two accessions differing at the *iucundus* locus and three Bryansk lines (Table S2-S4). PacBio libraries (one per genotype and organ) produced 14.2–22.3 M subreads for leaves (33.1–57.0 Gb) and 14.6–77.4 M (33.7–160.8 Gb) for stem. After SQANTI3 filtering, the clustered Iso-Seq transcriptome comprised 19,071 genes and 100,203 transcripts (N50 2,839 bp), increasing to 38,236 genes and 119,368 transcripts after adding gene models lifted from the NLL reference [31] to recover loci not captured by the long read data. Illumina libraries from leaf and stem yielded 82.0–102.9 million reads per genotype library (12.3–15.4 Gb per sample), with more than 97% of bases above Q20. Functional annotation assigned at least one GO term to 30,089 of the 38,236 genes (78.7%; Table S5). Relative to this, a merged transcriptome based on the previously published genome annotations v1 [32] and v2 [31] comprises 39,130 genes/transcripts (N50 1,701 bp).

### 4. Characterization of differentially expressed genes (DEGs) and candidate genes selection

Across all analyses, 18,851 genes met the significance threshold (Table S6). Separate DEG sets were generated for leaf and stem in *iuc* vs *Iuc* and Bryansk vs *Iuc* contrasts (Table S7-S10). The high confidence consensus DEG set comprised genes identified by all three methods (see Materials and Methods) and was used for subsequent analyses. In both contrasts leaf contributed more consensus DEGs than stem, comprising 663 DEGs in leaf (Table S9) and 315 in stem (Table S10) in *iuc* vs *Iuc*, and 622 in leaf (Table S7) and 352 in stem (Table S8) in the Bryansk vs *Iuc*.

Gene Ontology enrichment of the consensus DEGs showed that induced or suppressed biological processes differed, depending on both contrast and organ (Fig. 3; Table S11). Induced processes were largely shared in leaf and stem of *iucundus* and the Bryansk lines compared with *Iucundus*, whereas the repressed processes differed between both contrasts. In leaves, *iucundus* line showed reduced defense and hormone responses, as well as decreased transmembrane transport and catabolic processes. Bryansk leaves instead were depleted in responses to other organisms and toxic substances as well as in xenobiotic and amino acid transport, as well as hormone related processes. In stems, down-regulation in *iucundus* mainly affected flavonoid and anthocyanin biosynthesis and auxin and purine transport, while in Bryansk lines it primarily involved late endosome to vacuole transport. Shared induced categories in both organs of *iucundus* and the Bryansk lines included hydrogen peroxide catabolism, response to oxidative stress, responses to host organisms and defense mechanisms of other organisms, cellular development, antibiotic metabolism, lysine biosynthesis via diaminopimelate, cytokinin responses and Golgi vesicle budding. Additional, organ or genotype specific enrichments included carbohydrate metabolism in *iucundus* leaves, defense and cold responses in *iucundus* stem, proteolysis and hormone responses in the Bryansk lines leaves, as well as regulation of transcription by RNA polymerase II in Bryansk stems.

**Figure 3.**
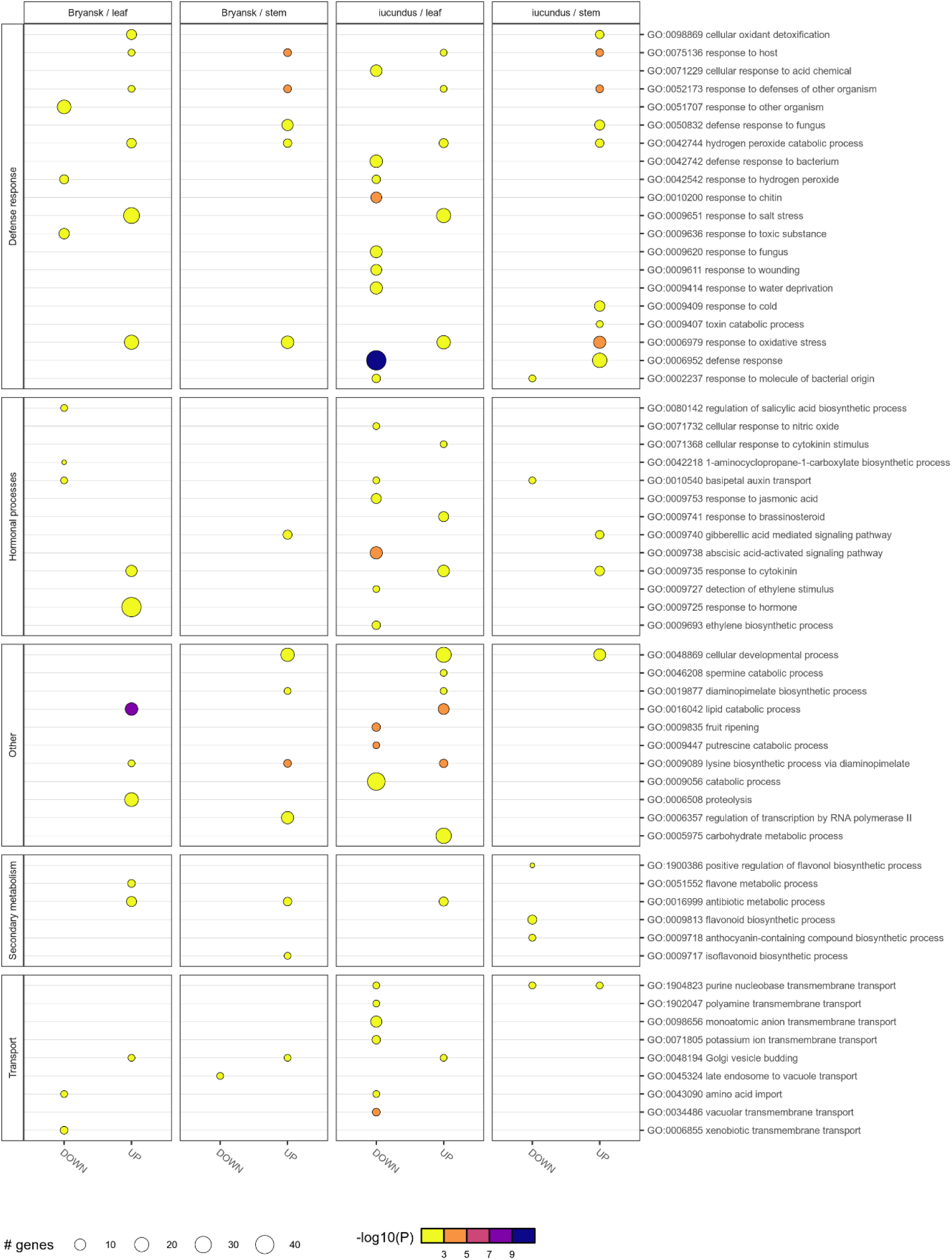
Gene ontology biological process enrichment for DEGs in leaf and stem of Bryansk lines and *iucundus* relative to *Iucundus*. Bubble plot showing enriched Gene Ontology biological process terms. Columns subdivide data by contrast/organ combinations with upregulated and down-regulated genes shown separately. Bubble size reflects the number of significant DEGs associated with each term and color indicates enrichment significance expressed on the minus log10 P-value scale.

To examine QA-related processes, further analyses focused on the down-regulated gene sets from both contrasts, comprising 259 and 96 genes in leaf and stem for the *iuc* vs *Iuc*, and 168 and 80 for the Bryansk vs *Iuc*. In total, 53 candidate genes were selected including 17 from the *iuc* vs *Iuc* contrast, 29 from the Bryansk vs *Iuc*, and seven known QA-related genes serving as a reference set (Table 2). For chromosomal context, candidate and known QA related genes from the *iuc* vs *Iuc* contrast were mapped onto the *iucundus* reference genome [31] (Fig. 4). This contrast was chosen for mapping because it combines candidate genes with coordinately down-regulated QA genes, including *RAP2-7* (PB.7870), and thus provides a clearer view of their chromosomal distribution. In *iuc* vs *Iuc*, candidates were mainly down-regulated in leaf but not stem, with fold change ranging from −23.40 to −5.52. In the reference set, *RAP2-7*, *LDC* (PB.16972), *LaCAO* (PB.16302), and *LaAT* (Lupan_033908) showed the same leaf restricted down-regulation. In contrast, *SDR1* (PB.16320) and *CYP71D189* (PB.3339), previously linked to *LDC* by co-expression [10], showed no consistent direction of change, and *HMT-HLT* (PB.10302) was not differentially expressed, consistent with prior reports [12]. In Bryansk vs *Iuc* contrast, all candidates were down-regulated in leaf and several in both organs, with fold change from −12.78 to −1.96. Across candidates, genes were distributed among overlapping functional categories, with most assigned to secondary metabolism, transcriptional regulation, and additional representation in transport and ion homeostasis, protein sorting and ethylene biosynthesis (Table 2).

**Figure 4.**
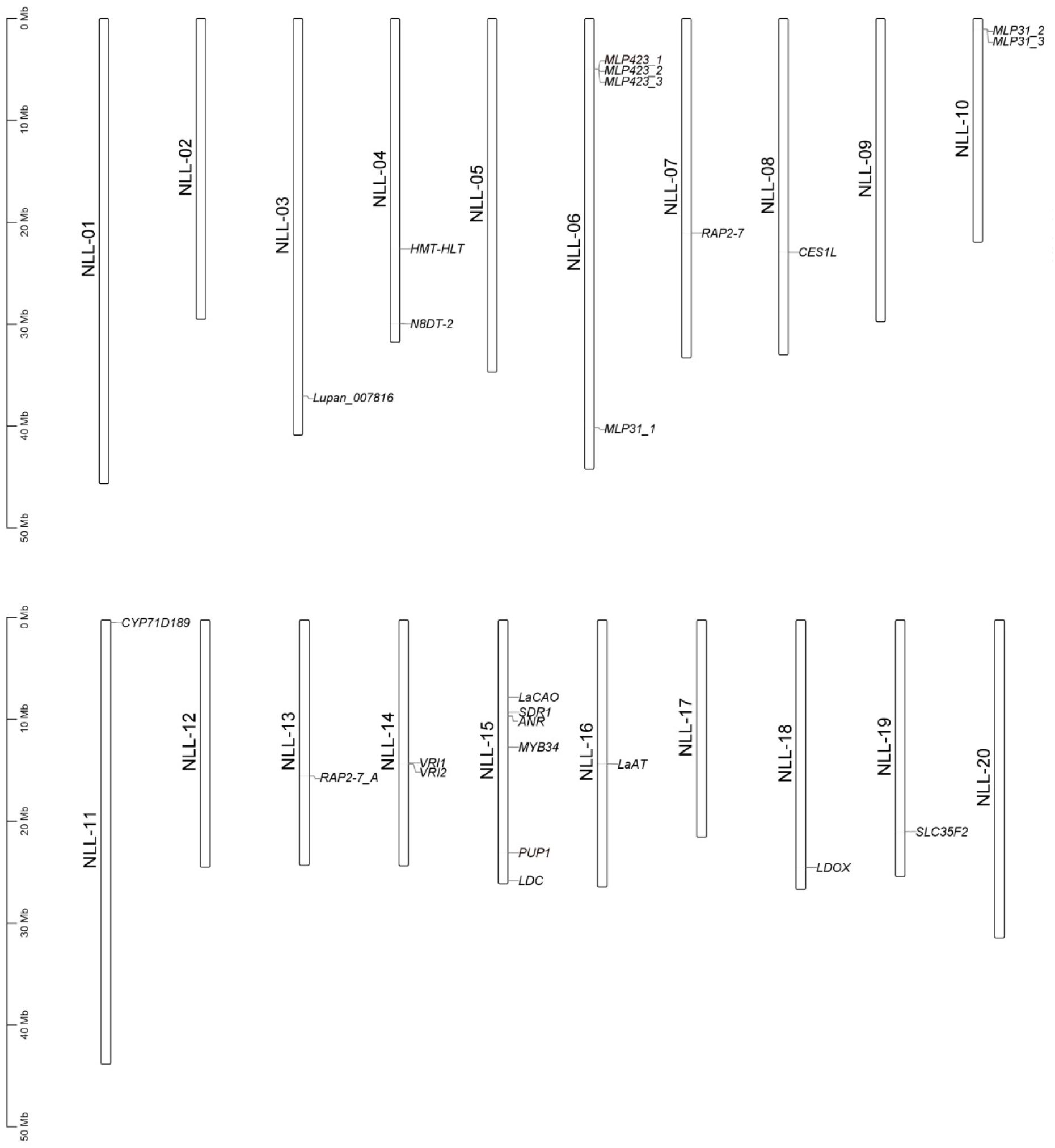
Chromosomal distribution of known QA pathway genes together with candidate genes from the *iucundus* versus *Iucundus* contrast in NLL. Map of the twenty NLL chromosomes in the *iucundus* reference genome [31] with chromosomes scaled in megabases and labels marking gene positions, generated using TBtools.

**Table 2.** Candidate genes involved in QA pathway with functional annotation and differential expression metrics in *iucundus* versus *Iucundus* and Bryansk versus *Iucundus* contrasts.

| Candidate gene set | Functional category | Gene ID (current) | Gene symbol | Gene IDs in previous annotations (v1, v2*) | Gene annotation (NCBI RefSeq, *Lupinus angustifolius*) | Linkage group (NLL) | DEG log2FC (E-value, method**) |  |  |  |
| --- | --- | --- | --- | --- | --- | --- | --- | --- | --- | --- |
|  |  |  |  |  |  |  | *iucundus* vs *iucundus*, leaf | *iucundus* vs *iucundus*, stem | Bryansk vs *iucundus*, leaf | Bryansk vs *iucundus*, stem |
| *iucundus* vs *iucundus* contrast | Secondary metabolism and biosynthesis | PB.16323 | **ANR** | TanjilG_14005, Lupan_020294 | anthocyanidin reductase ((2S)-flavan-3-ol-forming)-like | NLL-15 | -12.64 (3.99e-19; S) | 0 | 0 | 0 |
|  |  | Lupan_007209 | **CES1L** | TanjilG_32669, Lupan_007209 | carboxylesterase 1-like | NLL-08 | -5.52 (1.10e-13; S) | 0 | 0 | 0 |
|  |  | PB.14959 | **LDOX** | TanjilG_00883, Lupan_021727 | leucoanthocyanidin dioxygenase-like isoform X2 | NLL-18 | -23.03 (8.75e-09; S) | -34.56 (2.06e-20; S) | 0 | 0 |
|  |  | PB.2970 | **MLP31_1** | TanjilG_01236, Lupan_002572 | MLP-like protein 31 | NLL-06 | -15.38 (7.42e-32; S) | 0 | 0 | 0 |
|  |  | PB.20574 | **MLP31_2** | TanjilG_26043, Lupan_028507 | MLP-like protein 31 | NLL-10 | -5.62 (1.45e-09; R) | 0 | 0 | 0 |
|  |  | Lupan_028512 | **MLP31_3** | TanjilG_19334, Lupan_028512 | MLP-like protein 31 | NLL-10 | -13.91 (2.28e-28; S) | 0 | 0 | 0 |
|  |  | PB.2036 | **MLP423_1** | TanjilG_15923, Lupan_000661 | MLP-like protein 423 isoform X1 | NLL-06 | -14.86 (3.11e-29; S) | 0 | 0 | 0 |
|  |  | PB.2037 | **MLP423_2** | TanjilG_15922, Lupan_000662 | MLP-like protein 423 isoform X2 | NLL-06 | -13.40 (8.40e-20; S) | 0 | 0 | 0 |
|  |  | PB.2039 | **MLP423_3** | TanjilG_15921, Lupan_000663 | MLP-like protein 423 | NLL-06 | -8.03 (1.88e-14; R) | 0 | 0 | 0 |
|  |  | Lupan_011464 | **N8DT-2** | TanjilG_21382, Lupan_011464 | naringenin 8-dimethylallyltransferase 2, chloroplastic-like isoform X4 | NLL-04 | -6.94 (3.35e-11; R) | 0 | -4.82 (9.50e-09; R) | 0 |
|  |  | PB.19347 | **VRI1** | TanjilG_05322, Lupan_030549 | vestitone reductase-like | NLL-14 | -7.26 (2.00e-12; S) | 0 | 0 | 0 |
|  |  | PB.19348 | **VRI2** | TanjilG_05321, Lupan_030550 | vestitone reductase-like | NLL-14 | -6.05 (3.10e-10; S) | 0 | 0 | 3.08 (2.15e-03; k) |
|  | Transcription and gene expression regulation | PB.16358 | **MYB34** | TanjilG_10046, Lupan_036882 | transcription factor MYB34 | NLL-15 | -8.76 (1.12e-05; S) | 0 | 0 | 0 |
|  |  | PB.20273 | **RAP2-7A** | TanjilG_14185, Lupan_023724 | ethylene-responsive transcription factor RAP2-7-like | NLL-13 | -10.02 (2.42e-13; S) | 0 | 0 | 0 |
|  | Metabolite transport | PB.16702 | **PUP1** | TanjilG_28431, Lupan_020878 | purine permease 1-like | NLL-15 | -12.05(1.54e-23; S) | -3.27 (4.02e-04; k) | 0 | 0 |
|  |  | PB.17620 | **SLC35F2** | TanjilG_27659, Lupan_028023 | solute carrier family 35 member F2-like | NLL-19 | -23.40 (3.23e-15; R) | 0 | 0 | 0 |
|  | Other | Lupan_007816 | **Lupan_007816** | TanjilG_08196, Lupan_007816 | uncharacterized protein LOC109342541 isoform X1 | NLL-03 | -9.12 (2.83e-08; S) | 0 | 0 | 0 |
| Bryansk vs *Lucundus* contrast | Protein sorting and organelle trafficking | PB.3562 | **BLOS1** | TanjilG_24054, Lupan_032439 | biogenesis of lysosome-related organelles complex 1 subunit 1-like isoform X1 | NLL-11 | 0 | 0 | -4.05 (1.68e-65; **k**) | -5.17 (1.29e-122; **k**) |
|  |  | Lupan_002219 | **CHMP1B** | TanjilG_05990, Lupan_002219 | ESCRT-related protein CHMP1B-like | NLL-06 | 0 | 0 | -12.24 (2.37e-88; **S**) | -11.90 (7.20e-83; **S**) |
|  |  | Lupan_032803 | **IST1** | TanjilG_15149, Lupan_032803 | IST1 homolog [Arachis ipaensis]*** | NLL-11 | -2.89 (1.98e-03; **k**) | 0 | -3.20 (3.12e-07; **k**) | -3.44 (2.52e-04; **R**) |
|  | Transport and ion homeostasis | PB.9831 | **CAT** | TanjilG_15405, Lupan_009525 | cationic amino acid transporter 7, chloroplastic-like | NLL-04 | -3.17 (4.42e-03; **k**) | -3.82 (5.95e-05; **k**) | -3.44 (9.91e-06; **k**) | -2.90 (2.04e-03; **k**) |
|  |  | PB.8060 | **CAX3** | TanjilG_24316, Lupan_013395 | vacuolar cation/proton exchanger 3-like | NLL-07 | -6.34 (8.33e-08; **S**) | 0 | -5.54 (1.67e-08; **S**) | 0 |
|  |  | Lupan_015411 | **CNGC1** | TanjilG_02509, Lupan_015411 | cyclic nucleotide-gated ion channel 1-like isoform X2 | NLL-02 | -2.90 (1.93e-03; **k**) | 0 | -2.47 (3.99e-03; **k**) | 0 |
|  |  | Lupan_018450 | **DTX16** | TanjilG_02143, Lupan_018450 | protein DETOXIFICATION 16-like isoform X2 | NLL-05 | 0 | -2.19 (5.04e-04; **k**) | -2.13 (8.26e-06; **k**) | -2.69 (8.57e-13; **k**) |
|  |  | Lupan_008507 | **DTX21** | TanjilG_00065, Lupan_008507 | protein DETOXIFICATION 21-like | NLL-03 | -5.06 (1.32e-07; **k**) | 0 | -5.17 (2.01e-12; **k**) | 0 |
|  |  | PB.21510 | **KEA18** | TanjilG_25011, Lupan_024380 | cation/H(+) antiporter 18-like | NLL-17 | -4.32 (1.98e-07; **R**) | -4.13 (1.21e-06; **R**) | -4.15 (2.08e-10; **R**) | -3.41 (3.95e-06; **k**) |
|  |  | PB.2596 | **NPF6.3** | TanjilG_32231, Lupan_001540 | protein NRT1/ PTR FAMILY 6.3-like | NLL-06 | -3.44 (1.14e-04; **S**) | 0 | -2.57 (8.99e-04; **R**) | 0 |
|  |  | PB.19119 | **WAT1** | TanjilG_12690, Lupan_030170 | WAT1-related protein At4g08300-like | NLL-14 | -4.29 (1.47e-08; **R**) | 0 | -3.04 (2.44e-05; **R**) | 0 |
|  |  | Lupan_007045 | **ZIFL1** | TanjilG_22335, Lupan_007045 | protein ZINC INDUCED FACILITATOR-LIKE 1-like isoform X5 | NLL-08 | -3.95 (1.02e-72; **S**) | 0 | -3.26 (3.22e-81; **k**) | 0 |
|  | Ethylene biosynthesis | PB.12627 | **ACO4** | TanjilG_24114, Lupan_026415 | 1-aminocyclopropane-1-carboxylate oxidase homolog 4-like | NLL-09 | -3.81 (3.56e-10; **k**) | 0 | -2.98 (1.23e-07; **k**) | 0 |
|  |  | PB.14361 | **ACS** | TanjilG_04889, Lupan_022502 | 1-aminocyclopropane-1-carboxylate synthase-like | NLL-18 | 0 | 0 | -4.77 (1.47e-08; **R**) | 0 |
|  | Transcription and gene expression regulation | PB.6812 | **AP2_41710** | TanjilG_26837, Lupan_019639 | AP2-like ethylene-responsive transcription factor At2g41710 | NLL-05 | 0 | 0 | -1.96 (1.28e-03; **R**) | 0 |
|  |  | PB.16404 | **bHLH137** | TanjilG_25129, Lupan_020419 | transcription factor bHLH137-like isoform X1 | NLL-15 | 0 | 0 | -2.55 (7.45e-04; **R**) | 0 |
|  |  | PB.9362 | **bZIP11** | TanjilG_23462, Lupan_006525 | bZIP transcription factor 11-like | NLL-08 | -5.21(1.27e-12; **R**) | 0 | -3.50 (6.49e-07; **k**) | 0 |
|  |  | PB.4338 | **IAA4** | TanjilG_09391, Lupan_031517 | auxin-induced protein 22B-like | NLL-11 | 0 | 0 | -11.15 (1.06e-41; **S**) | -12.78 (2.10e-56; **S**) |
|  |  | PB.7173 | **MYB59** | TanjilG_03285, Lupan_014873 | transcription factor MYB59-like isoform X1 | NLL-07 | -3.25 (2.08e-04; **k**) | 0 | -2.60 (2.19e-03; **R**) | 0 |
|  |  | Lupan_024589 | **MYB108** | TanjilG_29228, Lupan_024589 | transcription factor MYB108-like isoform X2 | NLL-17 | -5.94 (7.47e-06; **k**) | 0 | -4.69 (1.80e-05; **k**) | 0 |
|  |  | PB.15930 | **RAP2.3** | TanjilG_27008, Lupan_033269 | ethylene-responsive transcription factor RAP2-3-like | NLL-16 | 0 | 0 | -3.14 (3.51e-05; **R**) | 0 |
|  | Secondary metabolism and biosynthesis | Lupan_010622 | **ARG2** | TanjilG_16486, Lupan_010622 | indole-3-acetic acid-induced protein ARG2-like | NLL-04 | -4.14 (1.04e-04; **k**) | 0 | -3.05 (3.84e-03; **k**) | 0 |
|  |  | Lupan_030034 | **ATL42** | TanjilG_12559, Lupan_030034 | E3 ubiquitin-protein ligase ATL42-like | NLL-14 | -2.47 (1.60e-05; **k**) | 0 | -2.54 (1.49e-09; **k**) | 0 |
|  |  | PB.17657 | **CES15** | TanjilG_00283, Lupan_028114 | probable carboxylesterase 15 | NLL-19 | -5.40 (1.09e-10; **k**) | 0 | -5.02 (8.82e-14; **k**) | 0 |
|  |  | Lupan_024256 | **CYP71A1** | TanjilG_05596, Lupan_024256 | cytochrome P450 71A1-like | NLL-17 | 0 | 0 | -7.59 (1.89e-32; **k**) | -8.35 (4.32e-28; **k**) |
|  |  | Lupan_005954 | **CYP81E8** | TanjilG_02640, Lupan_005954 | cytochrome P450 81E8-like | NLL-08 | -5.81 (3.01e-03; **k**) | -5.40(1.82e-04; **k**) | -9.25 (2.44e-05; **S**) | -4.12 (1.43e-03; **k**) |
|  |  | PB.14161 | **CYP82A4** | TanjilG_22427, Lupan_015682 | cytochrome P450 82A4-like | NLL-02 | -3.11 (1.89e-05; **R**) | 0 | -3.09 (7.90e-09; **R**) | 0 |
|  |  | Lupan_024909 | **7DLGT** | TanjilG_25542, Lupan_024909 | 7-deoxyloganetic acid glucosyltransferase-like | NLL-17 | -8.80 (9.58e-15; **R**) | -5.95(5.89e-22; **R**) | -9.58 (3.04e-40; **R**) | -7.48 (8.07e-51; **R**) |
|  |  | PB.19771 | **MXMT1** | TanjilG_17310, Lupan_023417 | 7-methylxanthosine synthase 1-like | NLL-13 | -3.86 (1.82e-08; **R**) | 0 | -3.49 (1.37e-11; **k**) | 0 |
| Known QA biosynthetic and regulatory genes | Secondary metabolism and biosynthesis | PB.3339 | **CYP71D189** | TanjilG_29685, Lupan_032119 | cytochrome P450 71D10-like | NLL-11 | 0 | 0 | 0 | 0 |
|  |  | PB.10302 | **HMT-HLT** | TanjilG_22250, Lupan_010746 | 13-hydroxylupanine O-tigloyltransferase-like | NLL-04 | 0 | 0 | 0 | 0 |
|  |  | Lupan_033908 | **LaAT** | TanjilG_21586, Lupan_033908 | spermidine coumaroyl-CoA acyltransferase-like | NLL-16 | -11.81 (1.95e-17; **S**) | 0 | 0 | 0 |
|  |  | PB.16302 | **LaCAO** | TanjilG_00530, Lupan_020242 | amine oxidase [copper-containing] zeta, peroxisomal-like [Gastrolobium bilobum]*** | NLL-15 | -16.15 (1.60e-35; **S**) | 0 | 0 | 0 |
|  |  | PB.16972 | **LDC** | TanjilG_09726, Lupan_020529 | ornithine decarboxylase-like isoform X1 | NLL-15 | -16.78 (1.30e-38; **S**) | 0 | 0 | 0 |
|  |  | PB.16320 | **SDR1** | TanjilG_00579, Lupan_020288 | (-)-isopiperitenol/(-)-carveol dehydrogenase, mitochondrial-like | NLL-15 | 0 | 0 | 0 | 0 |
|  | Transcription and gene expression regulation | PB.7870 | **RAP2-7** | TanjilG_07628, Lupan_013780 | ethylene-responsive transcription factor RAP2-7-like | NLL-07 | -4.25 (1.69e-05; **S**) | 0 | 0 | 0 |

\* v1 IDs are TanjilG\_XXXX from the NLL genome annotation of Hane et al. 2017 [32]; v2 IDs are Lupan\_XXXXX from the NLL genome annotation of Garg et al. 2022 [31].
\*\* R – RSEM, S – StringTie, k – kallisto refer to the differential expression method used.
\*\*\*Best hits reported from RefSeq Viridiplantae.

### 5. Variant profiling in candidate and known QA-related genes

Variant discovery was performed using RNA-seq dataset. In the *iuc* vs *Iuc* contrast variant analysis was focused on candidate genes identified in this contrast and known QA-related genes. In the Bryansk lines the analysis was restricted to the known QA-related genes. Quality controlled reads mapped to the reference genome yielded 471 variants in these loci, including SNPs and small insertion–deletion variants (InDels) (Table S12). Variants were assigned to impact categories (see Materials and Methods), and subsequent analyses considered only homozygous variants with moderate or high predicted impact. In total, 17 such variants were found across 9 candidate genes (with one to three variants per gene), whereas six variants were detected in three known QA-related genes.

### 6. Mapping of RAP2-7 TFBS

DAP-seq was used to profile RAP2-7 binding [11,12]. All libraries passed initial quality control, with fractions of reads in peaks (FRiP) in 7,8-13,6% range for all samples, however upon greenscreen filtering of artifactual signals, lower final FRiP estimates of 0.1-1.5% could be seen [33], reducing the number of peaks from 2158 to 1130 (Table S13). Across genotypes, the number of inferred TFBS varied, with 15 sites in 83A:476 and 464 in P27255, compared with 54, 232 and 537 sites in 95826, 95927 and 95928, respectively.

Inferred RAP2-7 DAP-seq TFBS were distributed across intergenic regions (66%), exons (11%), introns (11%), promoters (7%) and transcription termination regions (5%). Among the intergenic sites, 62% were positioned more than 10 kb upstream/downstream of the nearest TSS. After excluding intergenic peaks, 383 sites were linked to their nearest gene based on promoter proximity, representing 370 unique genes, 256 of which had biological function annotation. Integration with RNA-seq showed that only 8,6% of the genes associated with TFBS were differentially expressed for at least one contrast, indicating that transcription factor (TF) occupancy alone did not predict the direction of expression change.

### 7. RAP2-7 regulatory motif reconstruction and docking simulation

Using conservative greenscreen peak filtering, *de novo* motif discovery in P27255 identified a centrally enriched AP2/ERF-like RAP2-7 binding motif, with the consensus sequence GTACGAGGTTT (top MEME result) (Fig. 5). Notably, no groups of centrally enriched motifs were recovered for 83A:476. In the Bryansk lines the same main motif was recovered with genotype dependent prevalence across peaks, corresponding to CentriMo motif group with up to 76% occurrence in 95826. On comparison with JASPAR2024 reference motif database, significant similarities to SMZ (AT3G54990), TOE2 (AT5G60120), and RAP2-7 (AT2G28550) binding site motifs from *A. thaliana* were detected (Fig. 5B) [34].

**Figure 5.**
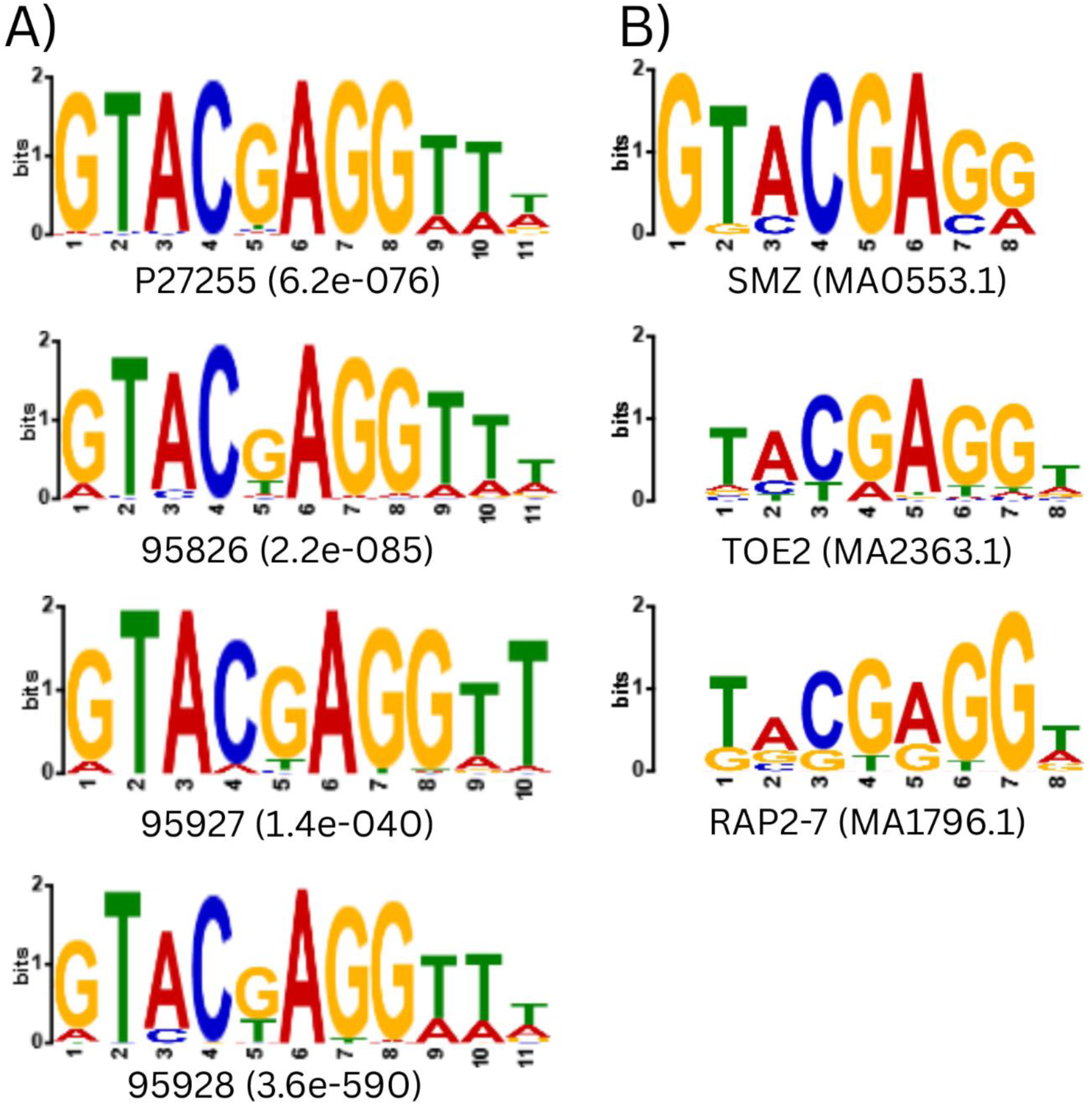
Comparison of RAP2-7 DAP-seq motifs across NLL genotypes and Arabidopsis homologs: A) sequence logos for RAP2-7 DAP-seq binding motifs in four NLL genotypes; B) known motifs of Arabidopsis SMZ, TOE2 and RAP2 7 retrieved from JASPAR (MA0553.1, MA2363.1, MA1796.1).

Previously, we have demonstrated that the arginine to serine substitution (R196S, relative to P27255 sequence) in NLL *RAP2-7* is strongly associated with low-alkaloid content [11]. To examine whether this mutation alters DNA motif recognition, we compared AlphaFold3 models of Tanjil RAP2-7, which carries the low-alkaloid S196 variant, and the alternative R196 version corresponding to high-alkaloid accessions, each bound to the reconstructed motif as DNA duplex. Although both structure models were of low quality (pTM=0.35 for the Tanjil reference and pTM=0.33 for the R196 variant), the AP2/ERF domain core (residues 149-211) was of considerably better quality (pDDT≥70). Prediction of hydrogen bonds based on these models indicates that R196 facilitates forming hydrogen bond with the second thymidine of the motif (dT2). Its additional contacts with adjacent residues helps stabilize the RAP2-7:DNA complex, whereas in the S196 variant this T2 centered interaction network is disrupted (see Fig. 6).

**Figure 6.**
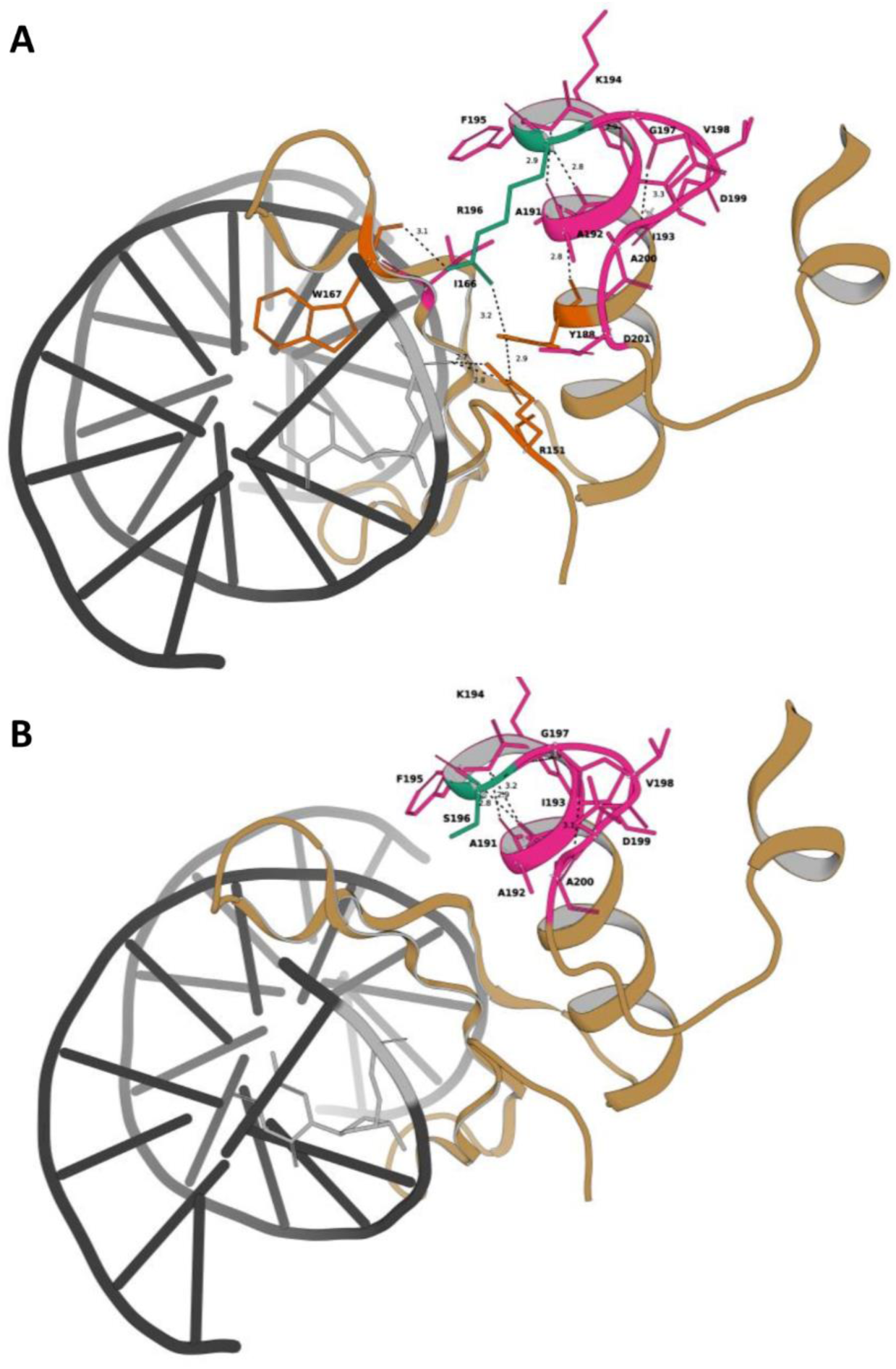
Residue R196 interactions in RAP2-7 variants in complex with DNA. (A) R196 in RAP2-7, representing the *Iucundus* allele, forms a hydrogen bond with dT2 of the motif. (B) S196 in RAP2-7, representing the *iucundus* allele, shows no clear network of hydrogen bonds to dT-2 of the binding motif. Residue 196 is shown in dark green. H-bonding partner residues in the vicinity of DNA are shown in orange, and additional R196-proximal protein residues interacting with its local environment are rendered in magenta. DNA is depicted in dark gray, with bases directly adjacent to the R196 microenvironment shown in a lighter gray. Distances marked over putative bonds are in angstroms (Å).

### 8. Integration of genome-wide RAP2-7 motif scanning with DAP-seq and differential expression

To better assess RAP2-7 binding, and to account for both DAP-seq sites that may have fallen below peak-calling thresholds and possible loss of binding competence due to the R196S substitution also present in the low-alkaloid NLL reference genome, cv. Tanjil [31], we used a two-part strategy. First, using the P27255 RAP2-7 reference motif, we compiled a comprehensive set of matching loci across NLL genome and cross-referenced these with differential expression results and DAP-seq regions. At a relaxed P-value threshold of 1×10⁻⁴, the scan returned 96 060 motif occurrences, 8 451 (8.8%) of which fell within 2 kb promoter regions. Of these, 684 occurred in association with DEGs (Table S14).

For the *iuc* vs *Iuc* DEGs, promoter-level contingency counts were: 515 DEGs with a motif, 37 721 DEGs without a motif, 7 312 non-DEGs with a motif, and 30 924 non-DEGs without a motif. Fisher’s exact test indicated strong depletion of the RAP2-7 motif among promoters of DEGs (odds ratio of approximately 0.058, P<2.2×10⁻¹⁶). For the Bryansk vs *Iuc* comparisons, promoter-level counts were: 527 DEGs with a motif, 37 709 DEGs without a motif, 7 300 non-DEGs with a motif, and 30 936 non-DEGs without a motif. Fisher’s exact test again indicated strong depletion of the RAP2-7 motif among promoters of DEGs (odds ratio of 0.059, P < 2.2×10⁻¹⁶).

Promoters of known and candidate QA-related genes (Table 2) were further inspected in this regard, yielding 14 putative TFBS across proximal regions of 12 DEGs (in 2 *iuc* vs *Iuc* candidates, 8 in Bryansk vs *Iuc* candidates, 2 supported by both types of contrasts) as well as non-differentially expressed *CYP71D189* known QA-related gene (Table 3). Strong matches were identified in particular in the promoters of the *RAP2-7* itself, the known QA gene *LaAT* and the candidate transcription factor *bZIP11* (PB.9362).

**Table 3.**
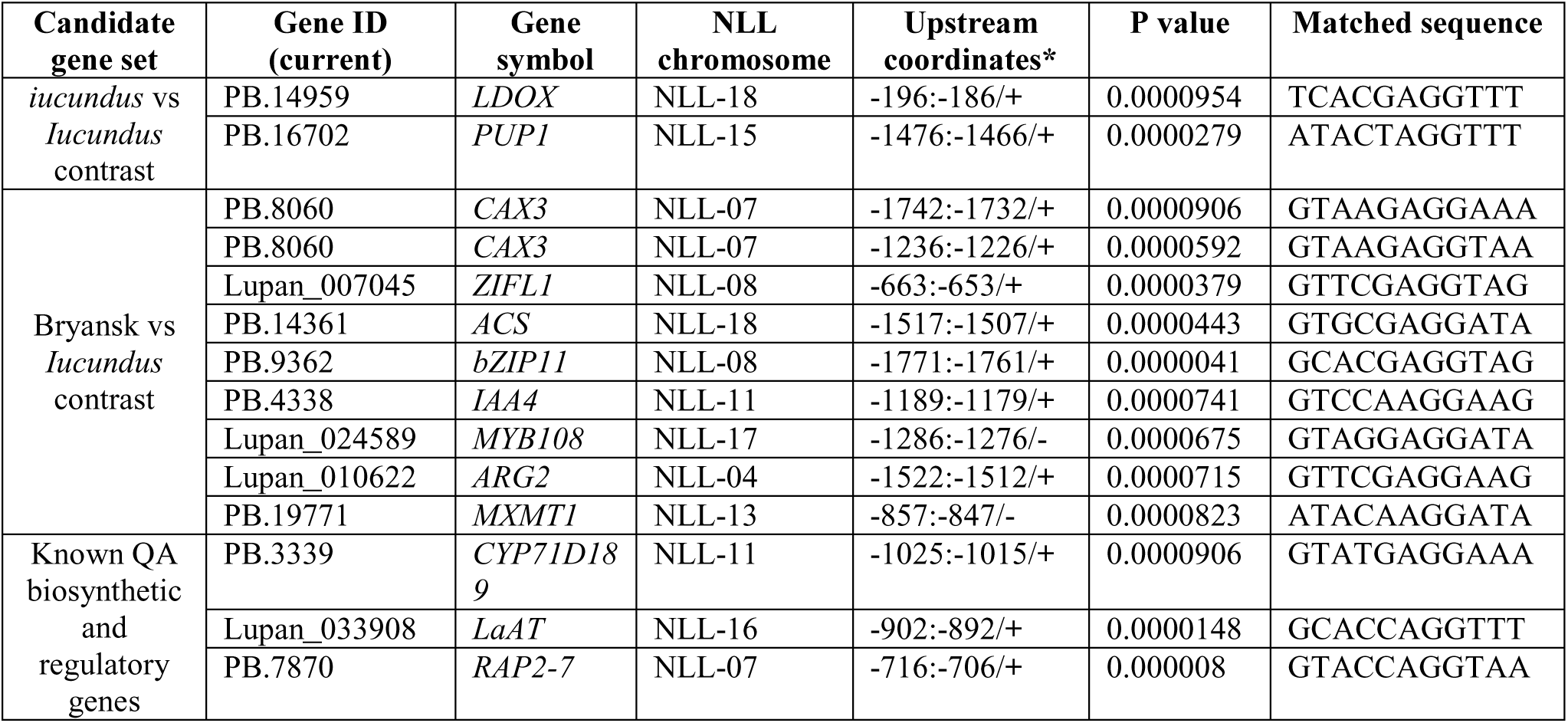
RAP2-7 binding motif occurrences in 2 kb promoters of candidate and known QA related genes. Full FIMO results for the NLL genome are provided in Table S14.

## Discussion

### Integrative QA pathway insights in *iucundus*-regulated genotypes

It has been shown that structural genes of the QA biosynthetic pathway display differential expression in leaves, stems and green pods in *iucundus* and *Iucundus* NLL genotypes [7,15,35]. Our qPCR and RNA-seq data for genotypes that differ at the *iucundus* locus show that the expression of *RAP2-7* and QA-related genes differ most strongly in leaves and pods and only weakly in stems (Fig. 2, Table 2). QA content in stems of 83A:476 remained low compared with P27255, but the fold difference between genotypes was smaller in stems than in either leaves or pods (Table 1). At organ level these patterns suggest that stems show only weak transcriptional regulation tied to QA differences under our experimental conditions, and may primarily contribute to transport and storage. However, QA biosynthesis in NLL has previously been traced to the epidermis of aerial organs, including stems [8], so such activity may have been masked in bulk stem samples. Together, our observations indicate, that leaf expression profiles provide the most informative basis for candidate gene selection, while a localized biosynthetic contribution of the stem epidermis cannot be excluded. Leaf DEG sets from contrasts that involve *iucundus* regulated genotypes encompassed known QA biosynthetic and regulatory genes, while simultaneously revealing new pathway candidates (Table 2). Notably, all candidates with expression data in the Lupin eFP Browser confirms elevated expression in the epidermis of QA producing leaves, stem, and pods, consistent with *LDC* and *LaCAO* profiles reported [8], which further supports their functional relevance.

Among the candidates, six loci encode major latex proteins (MLPs), members of the pathogenesis-related protein 10 (PR-10; Table 3). Four *MLP* genes map to NLL-06, forming a tight ∼12 kb cluster (*MLP423_1*/PB.2036, *MLP423_2*/PB.2037, *MLP423_3*/PB.2039), while the remaining two candidates co-localize on NLL-10 (Table 2, Fig. 4). Notably, although first linked to plant defense, MLP/PR10 proteins have also been recognized as enzymes in specialized metabolism, catalyzing reactions once considered spontaneous. Examples include: norcoclaurine synthase, thebaine synthase, and neopinone isomerase which mediate condensation, isomerization, and cyclization in BIA (morphine) biosynthesis [36–39]. These catalytic findings may have implications also for the QA pathway, particularly in light of the enzyme requirements recently proposed by Mancinotti and coauthors [9]. The authors proposed that tautomerization of Δ¹- to Δ²-piperideine, its stereoselective dimerization to tetrahydroanabasine, and analogous coupling to a quinolizidine intermediate yielding di-iminium cation are likely enzyme-catalyzed steps, that although chemically feasible without enzymes, ensure stereochemical control and pathway fidelity. Beyond catalysis, relevant opium poppy studies indicate that some MLP/PR10 proteins bind alkaloids, with latex-localized family members associating with specific BIAs and potentially contributing to their storage or transport [40]. Given that several QA biosynthetic, transport and storage steps still lack defined molecular components, our data supports MLPs as plausible mediators of some of these functions. *MLP* genes have previously been linked to QA accumulation in *Lupinus* and *Sophora* [35,41], but recent data further strengthen their role in alkaloid metabolism.

We identified two vestitone reductase-like (*VR1*/PB.19347 *VR2*/PB.19348) genes and one anthocyanidin reductase-like gene putatively forming (2S)-flavan-3-ols (*ANR*/PB.16323) (Table 2). Domain annotation classifies them as NAD(P) dependent dehydratase-like enzymes (IPR050425), placing them within the broader class of NAD(P)H dependent oxidoreductases. In a recently proposed QA pathway model, the tetracyclic di-iminium intermediate is converted to sparteine through two successive reductions attributed to one or more as yet unidentified reductases [9]. Given their predicted NAD(P)H dependent oxidoreductase activity and QA-like expression profiles *VR1-2* and *ANR* genes, are plausible candidates for these unresolved reductive steps. In this case, analogous NAD(P)H dependent reductions of cyclic iminium substrates are known from other alkaloid pathways. The noroxomaritidine reductase protein, employed during Amaryllidaceae alkaloid biosynthesis, also adopts the Rossman fold characteristic of NAD(P)H dependent oxidoreductases and belongs to the short chain dehydrogenase reductase (SDR) family [42]. In NLL, downstream conversion of sparteine to lupanine involves stepwise oxidation by SDR1 and CYP71D189 [10], underscoring the importance of enzyme mediating redox transformations within the QA pathway. Interestingly, although *VR1-2* and *ANR* were annotated as flavonoid-related, such *in silico* classifications may underestimate their metabolic relevance, and hide broader functional plasticity. This is also illustrated by tobacco (*Nicotiana tabacum* L.) A622 gene, originally classified as isoflavone reductase-like but required for pyridine alkaloid biosynthesis [43,44], and by dual ANR activity reported in proanthocyanidin biosynthesis [45]. Notably, *VRI1* and *VRI2* map to NLL-14 region about 105 kb diameter. *ANR* is located on NLL-15, within a ∼5 Mb interval that also harbors *La*C*AO*, *SDR1* and the candidate regulator *MYB34*, strengthening the candidacy of this locus for QA biosynthesis (Fig. 4). Together with their joint epidermal-enriched expression in QA-producing organs, this also supports other flavonoid-annotated candidates, including *LDOX* and *N8DT-2*, as potential contributors to QA biosynthesis.

Building on our previous identification of *RAP2-7* as a candidate regulator of QA biosynthesis [12], we now highlight a second *RAP2-7*-like transcription factor on NLL-13, hereafter referred to as *RAP2-7A* (PB.20273, TanjilG_14185; Fig. 4). *RAP2-7A* was previously reported as differentially expressed [12] and shows an even larger fold change in our current data. Moreover, we detected evidence that *RAP2-7A* may undergo premature termination in *iucundus* (Table S12), a phenomenon that could further contribute to the observed phenotype. Together with *MYB34*, this finding possibly expands the set of potential regulators involved in QA pathway regulation.

Of the other members of candidate set, *CES1L* (Lupan_007209) identified here and previously linked to a stable QTL for isolupanine [12] stands out as promising lead (Table 2). Carboxylesterase-like enzymes contribute to specialized metabolism, including BIA and MIA biosynthesis, where they mediate unconventional catalysis and ring forming cyclization [46,47]. In *iucundus*, *CES1L* carries three missense mutations, including a serine to proline change at position 256 (Table S12), and proline substitution in the catalytic domain which could alter active site architecture, enantioselectivity and substrate preferences [48]. Taken together, these combined features link *CES1L* to QA metabolism.

Last but not least, we identified several candidate transporters (Table 2). First, *PUP1* (PB.16702), an alkaloid transporter implicated in translocation between biosynthetic and storage sites, maps to NLL-15 (Fig. 4), the chromosome that carries several known and candidate QA genes, where it lies close to *LDC*. Although *PUP1*-like transporters have been reported in lupins, the specific role of this candidate in alkaloid-related processes remains unexplored [14]. In other species, related transporters mediate alkaloid movement and pathway control, for example tobacco NUP1 promotes nicotine uptake and also regulates ERF189, a key transcription factor in its pathway [49], while in opium poppy, BUP1 transfers alkaloids from phloem to laticifers for storage [26]. Second, we identified *SLC35F2* (PB.17620) a representative of the solute carrier transporter family conserved across eukaryotes, whose members mediate micronutrient uptake and drug transport in non-plant systems (e.g., yeast, protozoa, human cells) [50,51]. While, this latter gene lacks expression data in the Lupin eFP Browser [8], it ranks among the top DEGs in our contrasts.

### Insights into factors influencing QA metabolism in Bryansk lines

In our previous work [11], Bryansk lines combining low seed TAC, an *Iucundus* type RAP2-7 allele, and high leaf expression of QA-related genes could not be explained by the canonical *iucundus* mechanism, where low alkaloids are linked to RAP2-7-dependent SNP associated with repression of the core QA biosynthetic genes. We therefore considered alternative explanations, including altered expression of unknown QA genes, structural gene mutations, or impaired leaf to seed QA transport. Organ level TAC and qPCR, supported by multi-omics dataset, confirmed reduced QA content despite an *Iucundus* type expression profile of known QA genes across all examined organs (Table 1, Fig. 1), suggesting that the core biosynthetic machinery is largely intact. However, GO enrichment shows far fewer down-regulated categories in Bryansk vs *Iucundus* than in *iucundus*, involving distinct processes and making Bryansk functionally closer to *Iucundus* (Fig. 3). In Bryansk lines leaves, genes annotated as xenobiotic and amino acid transporters as well as those implicated in response to toxic substances are downregulated. This, taken together with down-regulation of late endosome to vacuole transport in stem samples, supports a sequestration bottleneck that may restrict leaf detoxification and stem vacuolar sequestration, contributing to low TAC across organs. Moreover, the Bryansk lines show a distinct QA chemotype than an intermediate state, with lower lupanine and higher sparteine, isolupanine, and oxolupanine than both references (Fig. 1, Table S1). Elevated levels of upstream precursor (sparteine), suggest a bottleneck in its conversion to lupanine with rapid downstream turnover. Notably, RNA-seq shows no directional changes in *CYP71D189* and *SDR1* expression (Table 2), although missense substitutions in the N-terminal region of CYP71D189 may affect the predicted signal peptide and adjacent region (Table S12). Other QA gene variants are few in number with low to moderate predicted impact (Table S12). Thus, coding disruption of known QA genes or leaf to seed transport defects alone is unlikely to explain low TAC. Instead, we posit that additional regulatory or metabolic constraints or reduced intracellular trafficking and detoxification capacity, could better guiding the selection of candidate genes for the Bryansk phenotype.

Consistent with the above, GO enrichment of late endosome to vacuole transport in stems and transcriptomic evidence indicates a potential disruption of the endosomal sorting complexes required for transport (ESCRT) in Bryansk lines (Table 2 and Fig. 3). These include genes encoding ESCRT-related proteins whose expression was down-regulated in Bryansk lines, including *CHMP1B* (Lupan_002219), *BLOS1* (PB.3562), and *IST1* (Lupan_032803). ESCRTs regulates cellular repertoires of receptors, transporters, and other membrane proteins [52]. Within endosomal transport, recognition of ubiquitinated cargo by ESCRT I initiates intraluminal vesicles formation, with BLOS1 facilitating cargo sorting. ESCRT III, including CHMP1 type proteins and IST1, then polymerizes to remodel membranes and mediate scission to form multivesicular bodies (MVB) [53,54]. Direct evidence that ESCRT-dependent MVB trafficking transports alkaloids is still lacking. However, Arabidopsis mutants with impaired vacuolar trafficking do show reduced vacuolar flavonoid accumulation, supporting a vesicle mediated contribution to the vacuolar sequestration of specialized metabolites [55]. Given that the vacuole is also the principal alkaloid sequestration site [56,57], altered ESCRT components such as CHMP1B and BLOS1 could influence QA accumulation in Bryansk lines.

The Bryansk lines additionally show reduced expression of several genes associated with hormonal processes (Fig. 3; Table 2), including ethylene biosynthesis, represented by *ACS* (PB.14361), and basipetal auxin transport, represented by *ZIFL1* (Lupan_007045). *ACS* acts as the rate limiting enzyme in ethylene biosynthesis and is tightly regulated by developmental and environmental cues, thereby controlling processes ranging from germination to defense responses [58,59]. *ZIFL1* belongs to a transporter family implicated in phytosiderophore efflux, auxin homeostasis, and proton antiport activity [60]. The effects of both ethylene and auxin on alkaloid biosynthesis appear to be species dependent. In *Nicotiana sylvestris* Speg. leaves, ethylene suppresses alkaloid accumulation, similarly, nicotine biosynthesis is reduced in tobacco roots upon auxin application [61]. In contrast, ethylene signaling enhances MIA biosynthesis in *C. roseus* [62]. Notably, there is also another auxin related candidate gene: *IAA4* (PB.4338; Table 2), which, among the genes highlighted here, stands out alongside *CHMP1B* as having no detectable transcript in the analyzed Bryansk samples (Table S6). In *Arabidopsis thaliana*, *IAA4* encodes an AUX/IAA repressor that is degraded in response to auxin, thereby releasing auxin response factors to activate downstream genes [63–65]. Beyond the reported suppressive effect of auxin on alkaloid production, AUX/IAA family members have also been shown to promote other specialized metabolic pathways, including glucosinolate biosynthesis in *Arabidopsis* and anthocyanin accumulation in apple (*Malus domestica*) [66,67]. Finally, it is worth to mention, that we identified a RAP2-7 binding motif in the promoter regions of *ACS*, *ZIFL1*, and *IAA4* (Table 3), suggesting a potential direct regulatory link, although its functional relevance remains to be established.

Other candidate genes with reduced expression in the Bryansk lines span functions from transcriptional regulation to secondary metabolism, as well as transport and ion homeostasis (Table 2). These include: DTX21 (Lupan_008507) and DTX16 (Lupan_018450), members of the multidrug and toxic compound extrusion (MATE) family of cation antiporters, whose involvement in vacuolar sequestration of endogenous alkaloids has been demonstrated in several species [56,57,68]. Bryansk candidates associated with responses to other organisms and toxic substances include also *CAX3* (PB.8060) and *ARG2* (Lupan_010622) (Fig. 3; Table 2), both of which contain a RAP2-7 binding motif in their promoter regions (Table 3). CAX3 contributes to intracellular pH homeostasis via cation proton exchange, thereby modulating the proton motive force required for MATE mediated transport [68,69]. *ARG2* encodes a small LEA5-like protein whose expression is auxin inducible [70]. In *Arabidopsis*, LEA5 family members have been implicated in regulating translation in mitochondria and chloroplasts [71], while chloroplasts are also a sites of LDC activity [3]. Among additional TFs, besides *IAA4* discussed above two other candidates stand out: *bZIP11* (PB.9362) and *MYB108* (Lupan_024589) (Table 2). Notably, both genes were also down-regulated in *iucundus* leaves, but their expression profiles differed from the previously seen pattern of the epidermis localized *iucundus* candidate genes [8]. Members of the bZIP and MYB families have been linked to the modulation of alkaloid biosynthesis across species. For example, MYBs such as *C. roseus* CrBPF1 and *Ophiorrhiza pumila* OpMYB1 affect strictosidine and camptothecin accumulation, and bZIPs such as *C. roseus* CrGBF1 and CrGBF2 act as repressors in indole-related pathways [72–74]. Among the genes discussed here, the specific connections of *CAX3*, *ARG2*, *bZIP11*, and *MYB108* to QA metabolism remain unclear. As for *ACS*, *ZIFL1*, and *IAA4*, the presence of a RAP2-7 binding motif in their promoters suggests a potential regulatory connection, making them promising targets for future studies.

### Towards a mechanistic view of RAP2-7 in QA biosynthesis

In the current study we combine allele specific DAP-seq, sequence variation and expression data to explore whether this association is reflected in the RAP2-7 binding landscape across contrasting regulatory backgrounds comprising *iucundus*, *Iucundus* and Bryansk lines. Comparative analysis of RAP2-7 DAP-seq profiles revealed a pronounced depletion of high confidence TFBS in *iucundus* compared with *Iucundus* and Bryansk lines (Table S13). In keeping with this, *de novo* motif discovery from the *Iucundus* and Bryansk peak summit regions recovered a strongly enriched RAP2-7 binding motif that is centrally positioned within peaks and closely matches the canonical recognition sequence defined for SMZ, TOE2 and RAP2-7 [34] (Fig. 5), whereas no centrally enriched motifs were recovered from *iucundus* peak regions. Notably, structure modelling and recovery of motif instances both point to cumulative, possibly parallel, influence of RAP2-7 on *LaAT* gene expression through RAP2-7 R196S substitution, as well as mutation of the binding site motif located within the gene promoter (GCACCAGGTTT, where the T2 motif position predicted to interact with R196 in high-alkaloid genotypes is itself altered).

The *Iucundus* RAP2-7 binding signature agrees with our previous phylogenetic analysis [12], which places RAP2-7 (TanjilG_07628) and RAP2-7A (TanjilG_14185) in a well-supported legume subclade sister to that containing Brassicaceae AP2 proteins SMZ and SNZ and closely related to TOE2, thereby filling a gap between candidate based genetic evidence and conserved AP2/ERF motif preferences. Promoter analysis supplemented DAP-seq findings, demonstrating matches to the recovered RAP2-7 motif present in and potentially affecting 13 candidate genes (Table 3). However, none of these loci exhibited RAP2-7 DAP-seq peaks and analysis of binding motif relationship with differential gene expression suggests depletion of predicted motif among DEGs as a whole. This suggests indirect action of RAP2-7 on a number of QA biosynthetic genes, either through intermediate factors (such as RAP2-7A, MYB34, MYB108 or bZIP11) or distal binding. It is also important to note that motif presence alone may not be sufficient to ensure detectable binding events under our assay conditions. This is consistent with large scale *A. thaliana* cistrome analyses and methodological assessments of DAP-seq, showing that sequence motif occurrences greatly exceed experimentally defined TFBS and that binding specificity is shaped by DNA context, motifs sharing within TF families, and chromatin’s and cofactors’ effects [29,30,75,76]. Consistent with studies showing that TF complexes can alter genomic binding profiles and motif usage [77,78], RAP2-7 binding *in vivo* may likewise depend on yet unidentified partners. In case of distal binding, a relevant precedent exists in form of the tomato GAME9, an AP2/ERF factor that controls glycoalkaloid biosynthesis via a MYC2–GAME9 complex bound to a distal enhancer within a defined chromatin domain [79]

Regardless of the above, our study provides the first DAP-seq dataset for lupin, defining an initial benchmark for species specific assay performance, including peak saturation and sensitivity. It is also important to consider, that all reads were mapped to the NLL reference genome Tanjil [31], which itself carries the *iucundus* allele, so *Iucundus* specific TFBS that have been weakened, lost or structurally altered in this background would be underrepresented, reducing both peak detection and apparent motif recovery. Similar patterns, where loss or weakening of a TFs mode of action is accompanied by decay and remodeling of TFBS in its corresponding regulatory elements, have been reported [80,81]. In addition to potential *cis* regulatory changes, the reduced number of high confidence peaks and the failure to reconstruct a clear, centrally enriched RAP2-7 motif observed for the *iucundus* allele are consistent with R196S substitution reducing RAP2-7 binding capacity. *In silico* modelling of RAP2-7 bound to DNA containing the reconstructed motif (Fig. 6) supports this view and together these observations provide a mechanistic insight into RAP2-7 polymorphism’s association with the low-alkaloid phenotype.

In summary, our results refine RAP2-7 from a positional candidate to a mechanistically supported, context dependent regulator of QA biosynthesis and implicate the R196S substitution as an important contributor to the low-alkaloid phenotype. At this stage, our findings also provide additional evidence supporting direct RAP2-7 action on several QA pathway genes, most notably *LaAT*, through binding of proximal promoter sequences.

## Conclusion

This study captured two distinct genetic architectures that shape QA regulation resulting in low alkaloid NLL phenotypes. These integrated analyses uncover candidate genes for previously missing steps in QA biosynthesis, reveal global expression changes associated with a non-*iucundus* mode of regulation, and link RAP2-7 binding specificity including the R196S substitution to the emergence of low-alkaloid phenotypes. As a whole our inquiry refines the working model of RAP2-7 function and provides a coherent set of testable hypotheses and genetic targets for future functional studies and rational improvement of low-alkaloid germplasm. The associated datasets also establish new, high quality transcriptomic resources for NLL, together with the first DAP-seq assay of NLL material, elucidating characteristics of RAP2-7 binding sites.

## Materials and Methods

### 1. Plant material and sample collection

Five NLL accessions were analyzed. Two carry contrasting alleles at the *iucundus* locus 83A:476 (*iucundus* allele, low-alkaloid, also referred to as sweet or *iuc*) and P27255 (*Iucundus* allele, high-alkaloid, also referred to as bitter or *Iuc*). The remaining three, 95826 (Bryanskij-35), 95927 (Bryanskij-123) and 95928 (Bryanskij-237/83) show an alternative mechanism underlying low-alkaloid phenotypes and are referred to as the Bryansk lines according to their origin [11]. In all datasets, each accession was analyzed individually. For interpretation and integrative comparisons across datasets, the three Bryansk lines were treated as a single group and compared with the *iucundus* and *Iucundus* regulatory types, collectively the references.

Plants were cultivated in a growth chamber (16 h photoperiod, 22/18 °C day/night, 55–60% relative humidity). Three independent cultivation experiments were conducted (2020, 2021, 2023), corresponding to successive stages of the work and covering all five accessions. Organs sampled for each assay block are detailed in Table 4. Samples collected for molecular analyses were flash frozen in liquid nitrogen and stored at −80 °C, while samples for alkaloid assessment were processed fresh.

**Table 4.** Organs sampled and assays performed for five NLL accessions (83A:476, P27255, 95826, 95927, 95928).

| Organ | qPCR, GC<sup>1</sup> | PacBio (Iso-Seq), Illumina (RNA-seq)<sup>2</sup> | DAP-seq<sup>3</sup> |
| --- | --- | --- | --- |
| Fully expanded young leaves | + | + | + |
| Stems | + | + |  |
| Flowers | + |  |  |
| Green pods (pod walls, without seeds) | + |  |  |
| Green, ripened seeds | + |  |  |
| Dry, ripened seeds | + |  |  |

<sup>1, 2, 3</sup> Each assay block was performed on organs collected within a single cultivation experiment.

### 2. Transcriptome sequencing

#### 2.1. RNA extraction

Total RNA was isolated from 30 mg of frozen, ground organ tissue (Table 4) using a Maxwell RSC Plant RNA Kit (Promega, Madison, WI, USA). RNA concentration and integrity were assessed with a NanoDrop spectrophotometer (Thermo Fisher Scientific, Waltham, MA, USA) and an Agilent 2100 Bioanalyzer (Agilent Technologies, Waldbronn, Germany), with a minimum RNA integrity number of 8. The same isolates were used to prepare PacBio Iso-seq and Illumina RNA-seq libraries.

#### 2.2. PacBio Iso Seq library construction and SMRT sequencing

Total RNA from five plants per accession was pooled in equal amounts to prepare a single Iso-Seq library (Table S15). Libraries were constructed following PacBio’s Iso-Seq sequencing protocol combining non-size selected and size selected fractions. Briefly, the first strand cDNA was synthesized using the SMARTer PCR cDNA Synthesis Kit (Clontech, CA, USA), amplified, and processed with the SMRTbell Template Prep Kit 1.0 (Pacific Biosciences, CA, USA). Sequencing was performed on the PacBio Sequel II system (HiFi/CCS mode; ≥20 GB raw data per sample) at Novogene Co. Ltd.

#### 2.3. Illumina RNA-Seq library construction and sequencing

cDNA libraries for Illumina RNA-seq were prepared separately for each replicate (Table S15). mRNA was purified with oligo(dT) magnetic beads, fragmented, and reverse-transcribed with random hexamer primers, followed by second-strand synthesis. Libraries were prepared through end repair, A-tailing, adaptor ligation, size selection, amplification, and purification. Sequencing was conducted on the Illumina NovaSeq platform (PE150) at Novogene Co. Ltd.

### 3. Transcriptome analysis

#### 3.1. Consolidated annotation of NLL reference

Merged reference annotation combined the draft Ensembl/Plants release 60 annotation [32] with the updated chromosome-length reference [31], accessed from LupinExpress on October 10 2024, with additional annotation from Prof. Lingling Gao (pers. comm.). Consensus annotation was generated with LiftOver v1.6.3 [82] using coverage ≥0.5, child feature identity of ≥0.9 (-s 0.9), CDS polishing (-polish), and CDS status flags (-cds), with mappings provided by minimap2 v2.28-r1209. The resulting GFF3 files were merged using gffutils, retaining features from the newer assembly and adding a qualifier with previous identifiers to overlapping records (Dataset S1).

#### 3.2. Reconstruction and curation of long-read transcriptome

Circular consensus sequences were generated using *pbccs* version 6.4.0 (minimum predicted accuracy 0.9). Primer removal and demultiplexing were performed using *lima* 2.13.0 in IsoSeq mode with peek-based primer inference. Full length non chimeric (FLNC) reads with detectable poly(A) tails were generated using *isoseq3 refine* (v4.0.0) and *pbtk* (v3.5.0), merged with *pbmerge*, standardized by assigning a unified SM tag in the BAM header, reheadered with SAMTOOLS (v1.21), and indexed with *pbindex*.

FLNC reads were clustered using *isoseq3 cluster2* (QV support), to generate a consensus BAM, aligned to the NLL reference genome with *pbmm2* (v1.17.0, ISOSEQ preset), and collapsed with *isoseq3 collapse* using clustered alignments plus the original FLNC BAM, preserving distinct 5′ exon structures.

The GFF file with collapsed high-confidence transcript models was curated using SQANTI3 v5.3.6 (*sqanti3_qc.py*) using all available Illumina reads, FLNC transcript counts and the post-LiftOver consensus annotation. Filtered models were then rescued using *sqanti3_rescue.py*. The full-splice matches (FSM) were required to have 0-59% adenine downstream of the transcription termination site. Non FSM transcript types were rescued if they met at least one of three criteria: (i) 0-59% downstream adenine, no reverse-transcription switching detected (RTS_stage=FALSE), isoform expression ≥50, and expression ratio ≥0.15; (ii) the same adenine and RTS_stage criteria plus fully canonical splice junctions; or (iii) the same adenine and RTS_stage criteria plus minimum coverage of three long reads. Transcripts meeting any criterion were retained, and reference models were added when no overlapping long-read derived model was present. The final annotation of [31] assembly in GFF3 format is included in the Supplementary Materials.

#### 3.3. Functional annotation

Functional annotation was performed with EnTAP v1.1.1 [83] searches using DIAMOND v2.1.8 [84] against NCBI/RefSeq Plants and UniProt/SwissProt (downloaded at 23/05/2024), and with eggnog-mapper v2.1.12 [85], using EggNOG 5.0 database [86] for ortholog inference. Annotations were manually curated to retain only general or plant-specific functional terms.

#### 3.4. Differential gene expression

Illumina RNA sequencing data were processed with the nf-core *rnaseq* pipeline v3.18 [87] (10.5281/zenodo.14537300) using the final annotation. Short reads were trimmed with TrimGalore v0.6.10 and cutadapt 4.9 at default settings, then mapped and quantified using three approaches: RSEM v1.3.1 [88] with STAR v2.7.10a [89], StringTie v2.2.3 [90] with STAR and kallisto v0.51 [91]. Outputs from all methods were imported into R with tximport v1.30.0, length normalized and modelled in DESeq2 v1.42, using a two-factor (genotype and tissue) generalized linear model with interaction. Likelihood ratios were performed by comparing full and reduced models. In pairwise comparisons, genes with baseMean ≥5, |log₂ fold change| ≥2 and adjusted P value ≤0.01 were reported as significantly differentially expressed. Gene Ontology (GO) enrichment was conducted with topGO v2.54 [92] using Fisher exact test (α=0.05) and parent-child weighting algorithm [93] to account for GO term hierarchy.

### 4. Candidate genes selection

Differential expression profiling and candidate genes selection were performed separately for the contrasts *iuc* vs *Iuc* and Bryansk vs *Iuc*, with the three Bryansk lines analyzed jointly as one group. The primary pool for candidate’s selection comprised genes significant in all three methods, with a small number manually incorporated from the broader set of significant DEG (significant according to at least one method). Final selection was based on functional annotations, GO term enrichment, expression in QA-producing organs [8], expression fold change, or prior literature evidence. This strategy yielded a robust list of candidate genes for each contrast and ensured that the final set included the high confidence consensus DEGs, while also capturing a few well supported candidates that might have been missed by strict filtering.

### 5. Variant calling of candidate and QA-related genes

Trimmed reads from nf-core/rnaseq pipeline were re-aligned with HISAT2 v2.2.1 [94] to the Tanjil reference in downstream-transcriptome–aware mode, providing splice junctions from the final annotation. Alignments were converted to sorted, duplicate marked BAMs with Sambamba v1.1.0 [95]. Variant calling followed Genome Analysis ToolKit v4.6.2 [96,97] best practices using AddOrReplaceReadGroups, SplitNCigarReads. and HaplotypeCaller with multithreaded pair-HMM and the --recover-dangling-heads option. Variants were annotated with Variant Effect Predictor v114.1 [98] to infer their impact on protein sequence and splicing sites. SignalP 5.0 [99] and Phobius [100] were used to assess whether variants detected in the N-terminal region of gene of interest fall within the predicted signal peptide.

### 6. *In vitro* Halo-Tag-RAP2-7 recombinant protein expression

#### 6.1. Constructing a Gateway Halo-Tag-RAP2-7 expression clones

Halo-Tag-RAP2-7 constructs were generated by Gateway recombination cloning technology and the pIX-Halo destination vector (ABRC stock CD3-1742) carrying N-terminal Halo-tag. RAP2-7 coding sequence was amplified using One-Phusion High-Fidelity DNA polymerase (GeneON GmbH, Groß-Rohrheim, Germany) and gene-specific primers (F: 5′-CACCATGGCTATGTTTGATCTCAAT-3′, R: 5′-TCAACTTGGGGAGTTAAATGAGGA-3′) with technical replicates per genotype, cloned into pENTR/SD/D-TOPO entry vector (Thermo Fisher Scientific) and sequenced (Genomed S.A., Warsaw) against the RAP2-7 reference sequence.

#### 6.2. *In vitro* synthesis of Halo-Tag-RAP2-7 recombinant protein

Halo-Tag-RAP2-7 was synthetized *in vitro* using the TNT T7 Coupled Reticulocyte Lysate System (Promega) in 50 µl reactions containing: 25 µl lysate, 2 µl buffer, 1 µl T7 RNA polymerase, 1 µl amino acid mixture lacking leucine and methionine (1 mM each), 1 µl RNasin RNase inhibitor (40 U µl⁻¹), 1 µl Mg(OAc)₂ (25 mM), and 1 µg of plasmid DNA template (verified Halo-Tag-RAP2-7 clones). Negative controls without plasmid DNA template were also included. Reactions were incubated for 2 h at 30 °C. Five microliters of lysate were used for Western blot analysis and the remaining volume was frozen at −80 °C for DAP-seq.

#### 6.3. Western blot analysis

Samples (25 µl) were denatured at 98 °C for 5 min, separated in PAAgels containing 10% acrylamide and electroblotted for 35 min onto a 0.22 µm nitrocellulose membrane (Schleicher & Schuell, Dassel, Germany) using a semi-dry blotting apparatus (Bio Rad, Hercules, CA, USA). Membranes were stained with Ponceau red to verify equal transfer, then incubated with Anti-HaloTag antibody (Promega; 1:1500) and HRP conjugated anti-mouse immunoglobulin G (Santa Cruz Biotechnology, Dallas, TX, USA; 1:5000) for 1 h each. Signals were detected with a WesternBright Quantum chemiluminescent substrate (Advansta, San Jose, CA, USA) and imaged using a G:BOX system (Syngene, Cambridge, UK). The expected size of Halo-Tag-RAP2-7 recombinant protein was ∼81 kDa. Analyses were carried out using protein extracts from 12 (83A:476, P27255) or 6 (the Bryansk lines) replications per genotype.

### 7. DAP-seq library construction and processing

#### 7.1. Libraries construction

DAP-seq libraries were prepared in three biological replicates per genotype.

High-molecular-weight genomic DNA for DAP-seq was extracted from frozen young leaves (3 g) using a phenol:chloroform:isoamyl alcohol protocol, and quantified fluorometrically using the Qubit dsDNA Broad Range Assay (Thermo Fisher Scientific).

Genomic DNA libraries were prepared as in [29], with minor modifications. Briefly, 7.5 µg gDNA in EB buffer (10 mM Tris-HCl, pH 8.5) was sonicated to ∼200 bp fragments (Bioruptor Plus, Diagenode), size-selected with AMPure XP beads (2:1 bead:DNA ratio; Beckman Coulter Life Sciences, Indianapolis, IN, USA), end repaired (Fast DNA End Repair Kit, Thermo Fisher Scientific) and purified (GeneJET PCR Purification Kit, Thermo Fisher Scientific). Next, samples were A-tailed (Klenow 3–5′exo-, New England Biolabs, Inc., Ipswich, MA, USA), purified again and ligated overnight with truncated Illumina TruSeq Y-adaptors (T4 DNA Ligase, New England Biolabs). Finally, libraries were cleaned with AMPure XP beads (1:1), eluted in 40 μl of EB, and quantified with a Qubit dsDNA Broad Range Assay (Thermo Fisher Scientific).

#### 7.2. DAP-seq procedure

Halo-Tag-RAP2-7 protein was immobilized on Magne HALO-Tag beads (Promega) (1 h, RT). Beads were washed five times with 1× PBS containing 0.005% NP 40 (PBS-NP40), and then incubated with 1 µg of adaptor-ligated DNA library for 60 min. Negative controls were processed in parallel, using adaptor ligated DNA libraries from each accession incubated with beads withouut recombinant protein. Protein–DNA complexes were washed eight times with PBS-NP40 and bound DNA was eluted in 25 μl EB at 98 °C for 10 min. DNA was amplified with Phusion High-Fidelity DNA Polymerase (NEB) using Illumina TruSeq dual indexed primers (Primer A: 5’AATGATACGGCGACCACCGAGATCTACAC-NNNNNNNN-ACACTCTTTCCCTAC and Primer B: 5’CAAGCAGAAGACGGCATACGAGAT-NNNNNNNN-GTGACTGGAGTTCAG, where NNNNNNNN represents the i5/i7 index sequences). Final libraries were purified with AMPure XP beads (1:1), quantified with Qubit dsDNA High Sensitivity assay, and sequenced on an Illumina NovaSeq X Plus (PE150, Novogene Co. Ltd.), yielding ∼6 Gb of raw data per sample (≥20 million reads per sample). Negative control libraries underwent the same procedure and were used to filter background peaks during peak calling.

#### 7.3. DAP-seq data analysis

DAP-seq data were processed with the nf-core chipseq pipeline v2.1.0 [87] (10.5281/zenodo.13899404), adapted to this assay. Short reads were trimmed with TrimGalore v0.6.7 and cutadapt v3.4, aligned with Bowtie2 v2.5.2 [101] and peaks called with MACS3 v3.0.1 [102] in narrow peaks mode (FDR 0.01, genome size 613189654, read length 150), restricting analyses to chromosome scaffolds. Peaks were annotated with HOMER v4.11 [103] *annotatePeaks.pl* script. The resulting peaks were filtered against a blacklist obtained from MACS3 reanalysis of all controls following the “greenscreen” procedure [33], as peak areas supported by at least half of the samples. Centrally enriched consensus motifs were identified with MEMECHIP v5.5.8 [104], against genomic sequence masked using RED, and compared to the JASPAR 2024 plants nonredundant TFBS subset [105].

To ascertain presence of additional RAP2-7 TFBS, not detected by DAP-seq, NLL genome was additionally investigated with FIMO v5.5.8 [106] at P-value threshold of 0.0001 and using background frequencies derived from RED-masked genome. Association between the distribution of RAP2-7 motif and differential gene expression was assessed with Fisher exact test as implemented in R (v4.3.1).

### 8. Structural modelling of RAP2-7:DNA complexes

Approximate models were created using AlphaFold3 [107] webserver with RAP2-7 protein sequences differing in alkaloid context associate polymorphism (S196R, based on Tanjil reference NCBI/RefSeq protein XP_019435537.1) [11] and homodimer DNA corresponding to the majority rule consensus of the 83A:476 predicted STREME motif (AGTACGAGGTTT). Structure visualization was done in PyMol (version 3.1.0) and hydrogen bonds predicted by HBPLUS [108].

### 9. Gene expression analyses (qPCR)

Total RNA was extracted as detailed in section 2.1, and 1 µg was reverse transcribed with the Transcriptor First Strand cDNA Synthesis Kit (Roche, Mannheim, Germany) using anchored oligo(dT)18 primers. For each accession, cDNA from two technical replicates was pooled for gene expression analysis.

Expression of four alkaloid-related genes was quantified: APETALA2/ethylene response (AP2/ERF) transcription factor (*RAP2-7*, TanjilG_07628) [12], lysine decarboxylase (*LDC*, TanjilG_09726) [3], copper amine oxidase (*LaCAO*, TanjilG_00530) [14], acyltransferase-like gene (*LaAT*, TanjilG_21586) [13]. Primers and probes followed [12] and data were normalized to *ADH3*, *TUBA*, and *ELF1B* identified as the most stable reference genes across lupin organs [7]. Reactions were run on a LightCycler® 480 II system (Roche, Mannheim, Germany) with LightCycler 480 Probes Master (Roche) and TaqMan probes (Genomed), in 10 µL reaction, with three biological replicates, two technical replicates and a no-template control. Relative expression was calculated with the efficiency corrected method [109], using accession 83A:476 as the calibrator, and PCR efficiencies for target and reference genes ranged from 0.95 to 1.0.

### 10. Alkaloid extraction and quantification

Quantitative and qualitative QA profiles were evaluated in three biological replicates per accession across six organs (Table 4). Alkaloid were measured by gas chromatography (GC-FID, GC-2014; Shimadzu, Kyoto, Japan), using the calibration equation y = 0.0159x, where x is the ratio of lupanine to caffeine peak areas and y is lupanine content, (mg). Extraction and analysis were performed as described previously [17,18]. Total alkaloid content (TAC) was expressed as % seed dry weight, and individual alkaloids were quantified by their relative abundance, as percentage of TAC (defined as 100%).

## Acknowledgements

This work was supported by the National Science Centre, Poland (MINIATURA 3, 2019/03/X/NZ1/02009), and the Polish Ministry of Agriculture and Rural Development under the Program of Basic Research for Biological Progress in Crop Production (Task 19, Journal of Laws 2020, item 2016). We thank Prof. Andrea Gallavotti and Mary Galli, M.Sc., from the Waksman Institute of Microbiology, Rutgers University, for their methodological guidance on the DAP-seq procedure.

## Author contributions

Conceptualization: M.K.; Supervision: M.K.; Investigation: K.C., M.K., A.K-M., A.B.; Formal Analysis: G.K.; Funding acquisition: M.K.; Writing—original draft preparation: K.C., M.K.; Writing—review & editing: K.C., M.K. A.K-M., G.K. K.S., A.B.; Visualization, G.K., K.C., M.K. All authors reviewed and approved the final submission.

## Data availability

All genomic and transcriptomic data, generated in this study are available at ArrayExpress (EMBL-EBI) under the following accession numbers: E-MTAB-16429 for PacBio data, E-MTAB-16431 for Illumina RNA sequencing data, and E-MTAB-16430 for DAP-seq data.

## Supplementary information

Supplementary data supporting this study (Tables S1-15 and Dataset S1) are available on Figshare (DOI: 10.6084/m9.figshare.30739052).

